# Safety, pharmacokinetics, and liver-stage *Plasmodium cynomolgi* effect of high-dose ivermectin and chloroquine in Rhesus Macaques

**DOI:** 10.1101/2020.04.27.065409

**Authors:** Pattaraporn Vanachayangkul, Rawiwan Im-erbsin, Anchalee Tungtaeng, Chanikarn Kodchakorn, Alison Roth, John Adams, Chaiyaporn Chaisatit, Piyaporn Saingam, Richard J. Sciotti, Gregory A. Reichard, Christina K. Nolan, Brandon S. Pybus, Chad C. Black, Luis A. Lugo, Matthew D. Wegner, Philip L. Smith, Mariusz Wojnarski, Brian A. Vesely, Kevin C. Kobylinski

## Abstract

Previously, ivermectin (1–10 mg/kg) was shown to inhibit liver-stage development of *Plasmodium berghei* in orally dosed mice. Here, ivermectin showed inhibition of the *in vitro* development of *Plasmodium cynomolgi* schizonts (IC_50_ = 10.42 μM) and hypnozoites (IC_50_ = 29.24 μM) in primary macaque hepatocytes when administered in high-dose prophylactically but not when administered in radical cure mode. The safety, pharmacokinetics, and efficacy of oral ivermectin (0.3, 0.6, and 1.2 mg/kg) with and without chloroquine (10 mg/kg) administered for seven consecutive days was evaluated for prophylaxis or radical cure of *Plasmodium cynomolgi* liver-stages in Rhesus macaques. No inhibition or delay to blood-stage *P. cynomolgi* parasitemia was observed at any ivermectin dose (0.3, 0.6, and 1.2 mg/kg). Ivermectin (0.6 and 1.2 mg/kg) and chloroquine (10 mg/kg) in combination were well-tolerated with no adverse events and no significant pharmacokinetic drug-drug interactions observed. Repeated daily ivermectin administration for seven days did not inhibit ivermectin bioavailability. It was recently demonstrated that both ivermectin and chloroquine inhibit replication of the novel Severe Acute Respiratory Syndrome Coronavirus 2 (SARS-CoV-2) *in vitro*. Further ivermectin and chloroquine trials in humans are warranted to evaluate their role in *Plasmodium vivax* control and as adjunctive therapies against COVID-19 infections.

## Introduction

Novel chemoprophylactic therapeutics and vector control interventions could support and accelerate malaria elimination efforts. Ivermectin mass drug administration (MDA) has been proposed as a malaria control tool since it makes the blood of treated persons lethal to *Anopheles* mosquitoes, the vectors of malaria (1–5), and repeated ivermectin MDAs in Burkina Faso were able to reduce malaria transmission to humans (6). Ivermectin is a safe and well-tolerated endectocidal drug used widely in veterinary and human medicine to combat both internal and external parasites.

Ivermectin has been shown to inhibit liver-stage development of *Plasmodium berghei* in both an *in vitro* Huh7 human hepatoma cell line model (7) and an *in vivo* C57BL/6 mouse model (8). The *in vitro* half maximal inhibitory concentration (IC_50_) for ivermectin *P. berghei* inhibition, IC_50_ = 1.8 µg/ml (2.1 µM), was higher than blood levels that can be achieved in treated humans. However, mice that were orally dosed with ivermectin at 1-10 mg/kg at 24 and 12 hours before and 12 hours after sporozoite challenge demonstrated liver-stage inhibition equal to primaquine (10 mg/kg) under the same dosing schedule (8). Human equivalent dosing (HED) that was evaluated in mice would correlate to ivermectin doses in the range of 0.08 – 0.81 mg/kg (9). Thus, ivermectin is promising for human malaria chemoprophylaxis as ivermectin doses as high as 2 mg/kg have been safely administered to humans (10). If ivermectin can prevent *Plasmodium* liver-stage infection, then ivermectin chemoprophylaxis could be considered in high risk groups such as forest-goers in the Greater Mekong Subregion or naïve soldiers deployed to malaria endemic areas. Furthermore, if ivermectin MDA is deployed for community-wide malaria vector control, and ivermectin is chemoprophylactic, then there would be direct benefits to MDA participants in preventing malaria infections.

*Plasmodium cynomolgi* infections in Rhesus macaques (*Macaca mulatta*) are routinely used as a surrogate human liver-stage model for *Plasmodium vivax* drug development. This model can evaluate both the causal prophylaxis, (*i.e.* protection from developing liver schizonts), and the hypnozoiticidal (*i.e.* radical cure of liver hypnozoites) efficacy of compounds (11). Ivermectin has been used in Rhesus macaque colonies to treat mites (12), lice (13), and intestinal helminths, such as *Ascaris*, *Trichuris*, and *Strongyloides fulleborni* (14–16). Pre-clinical studies demonstrated that oral ivermectin was safe in macaques at doses up to 1.2 mg/kg for 14 days and that macaques are an ideal animal model for ivermectin human treatment (17, 18). However, no study to date has evaluated the pharmacokinetics of repeated ivermectin treatment in Rhesus macaques or in combination with chloroquine.

Here we evaluate the *in vitro* and *in vivo* liver-stage effect of ivermectin against *P. cynomolgi* in Rhesus macaque liver hepatocytes and infected macaques. The safety and pharmacokinetics of repeated oral ivermectin dosing with and without chloroquine in macaques is also presented.

## Results

### In vitro results

Ivermectin efficacy against liver-stage parasites was initially evaluated using an *in vitro P. cynomolgi* liver model which utilizes primary Rhesus macaque hepatocytes in order to closely resemble the *in vivo* anti-relapse mode. The drugging regimen was defined by treatment mode, either prophylactic mode (*i.e.* drug administered with sporozoites and 3 days thereafter) or radical cure mode (*i.e.* drug administered from days 4 to 7 post sporozoite infection) similar to previously described methods (19). In prophylactic mode, ivermectin showed marginal *in vitro* causal protection against the development of *P. cynomolgi*-infected rhesus macaque hepatocyte liver schizonts IC_50_ = 9.12 μg/ml (10.42 μM) and hypnozoites IC_50_ = 25.59 μg/ml (29.24 μM) (Figure 1). However, in radical cure mode, ivermectin had no activity on developing *P. cynomolgi* liver schizonts or established hypnozoites, even when dosed at a high initial concentration of 100 µg/ml (114.26 μM).

**Figure 1).**
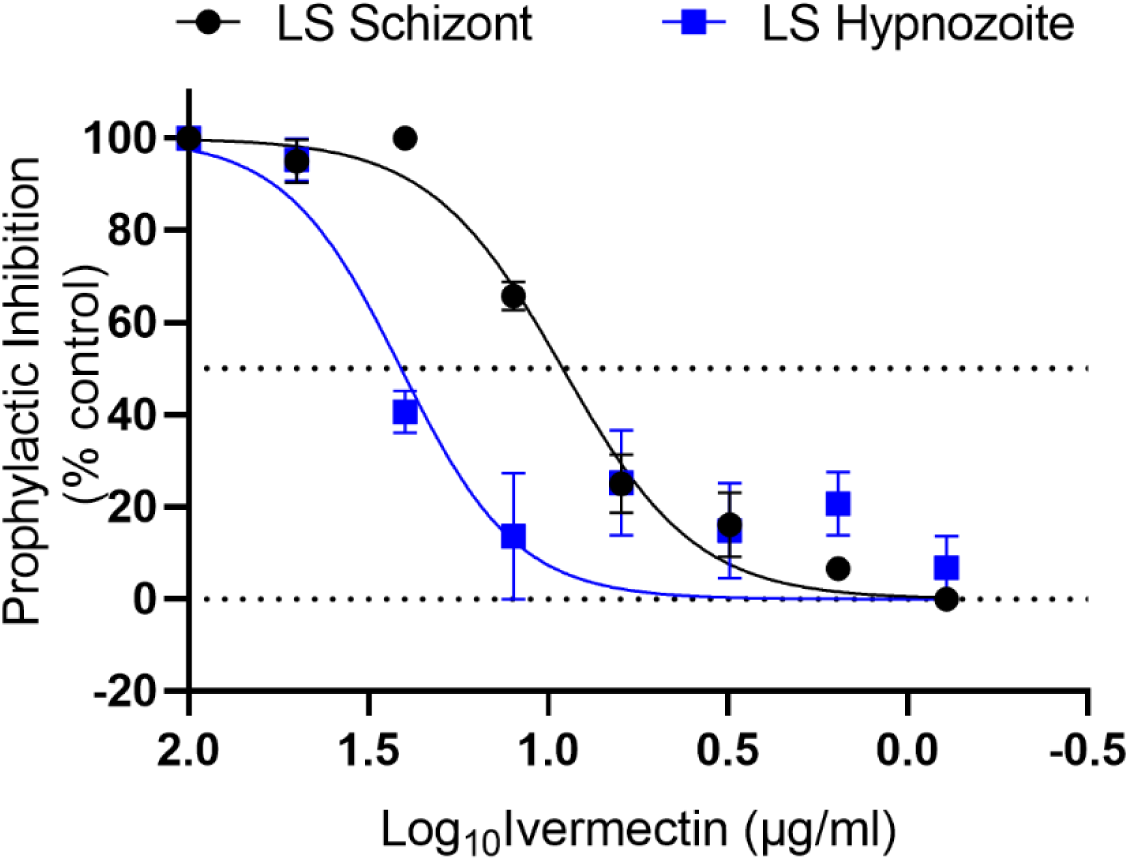
*In vitro Plasmodium cynomolgi* liver-stage ivermectin inhibition prophylactic results. Prophylactic (days 1-3) exposure of *P. cynomolgi* to ivermectin demonstrated marginal inhibition of liver schizonts (IC_50_ = 9.12 μg/ml) and hypnozoites (IC_50_ = 25.59 μg/ml). LS = liver-stage. Graph bars represent means with standard deviation of biological replicates (n = 3) with experimental replicates (n = 2).

### In vivo results: Ivermectin and chloroquine safety and tolerability

There was only one adverse event in a single macaque (R1435) that vomited three hours after the first oral dose of ivermectin (1.2 mg/kg) when administered as monotherapy one day prior to *P. cynomolgi* sporozoite injection. No adverse events occurred when ivermectin (0.6 or 1.2 mg/kg) was co-administered with chloroquine. No abnormal hematology outcomes were observed for ivermectin alone or ivermectin plus chloroquine co-administration.

### In vivo results: Parasitemia

Primary blood-stage parasitemia greater than 5,000/μl was detected ten days post inoculation for negative and positive control groups and for 2 of 3 macaques in both ivermectin high (1.2 mg/kg)- and low (0.3 mg/kg)-dose groups, with remaining macaques from each group reaching greater than 5,000/µl eleven days post inoculation which was 5 and 6 days after the last ivermectin administration. Primary infection blood-stage parasitemia was cleared from the negative control group with ten days of chloroquine (10 mg/kg) and both blood- and liver-stage parasites from positive control group with seven days of chloroquine (10 mg/kg) and primaquine (1.78 mg/kg). Blood-stage parasitemia was cleared from the three macaques in the low-dose ivermectin group with seven days ivermectin (0.6 mg/kg) and ten days chloroquine (10 mg/kg). Two of three macaques were cleared of primary infection blood-stage parasitemia in the high-dose group with ivermectin (1.2 mg/kg) for seven days and chloroquine (10 mg/kg) for ten days, while one macaque was cleared with ivermectin (1.2 mg/kg) and chloroquine (10 mg/kg) for seven days. However, the first relapse occurred within 3 weeks, at approximately the same time for negative control and both ivermectin groups with no significant differences for time to blood-stage parasitemia or treatment (Log-Rank (Mantel Cox) test P > 0.05). The first relapse infection blood-stage parasitemia was cleared from the negative control with chloroquine (10 mg/kg) alone for seven days. First relapse infection blood-stage parasitemia was cleared from both high (1.2 mg/kg)- and low (1.2 mg/kg)-dose ivermectin groups when given in combination with chloroquine (10 mg/kg) for seven days. Approximately 3 weeks later, a second relapse occurred in all negative control and ivermectin high- and low-dose treated macaques with no significant differences for time to blood-stage parasitemia or treatment (Log-Rank (Mantel Cox) test P > 0.05). At the point of second relapse, all ivermectin-group macaques were treated with primaquine (1.78 mg/kg) and chloroquine (10 mg/kg) for seven days. The positive control group was treated with primaquine (1.78 mg/kg) and chloroquine (10 mg/kg) for seven days at point of primary infection and had no relapses for the remainder of the study (Figure 2). The negative control group was treated with primaquine (1.78 mg/kg) and chloroquine (10 mg/kg) for seven days at the point of third relapse (data not shown).

**Figure 2).**
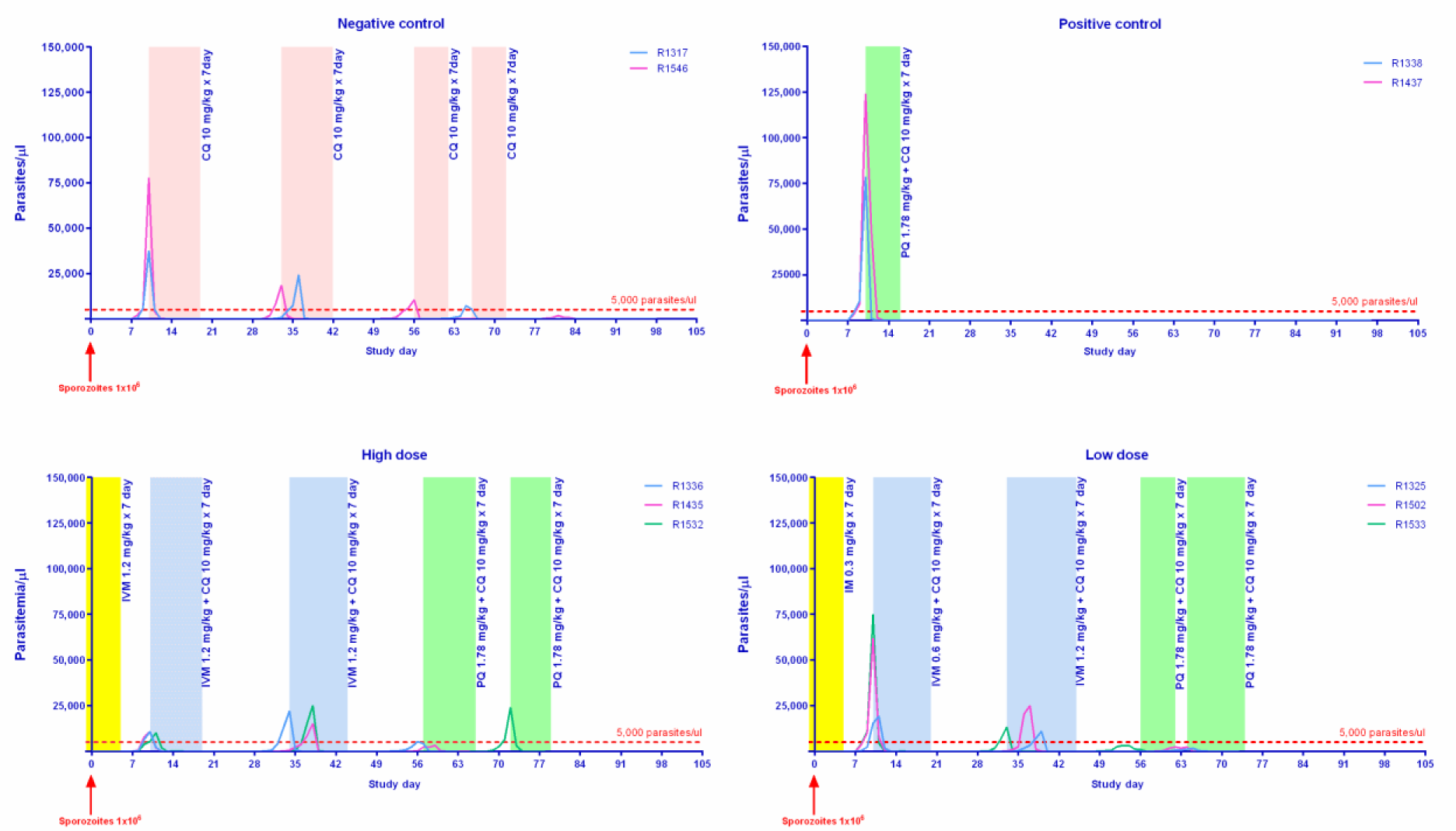
Blood-stage *Plasmodium cynomolgi* parasitemia results and drug regimen for each treatment group

Figure 2 displays the number of *P. cynomolgi* blood-stage parasites per μl of blood. Shaded areas represent the duration of drug administration when daily drug dosing was administered: yellow for ivermectin, peach for chloroquine, blue for ivermectin plus chloroquine, and green for primaquine plus chloroquine. Numbers in the legend denote the individual macaque identification number. The dashed red line denotes the 5,000 parasites per µl cutoff to trigger drug administration. IVM = ivermectin, CQ = chloroquine, PQ = primaquine.

The qRT-PCR method detected primary blood-stage parasitemia one day earlier than microscopy at point of first infection for the negative and positive control group macaques and in two out of three ivermectin low-dose (0.3 mg/kg) macaques. The remaining four ivermectin high- and low-dose macaques had blood-stage parasitemia detected by qRT-PCR on the same day as microscopy.

### In vivo results: Pharmacokinetics

Plasma ivermectin with and without co-administration of 10 mg/kg chloroquine reached maximum concentration (C_max_) at approximately 2-4 hours post-dose and the elimination half-life ranged from 11-28 hr with accumulation index of 0.6-3.7. Plasma concentration time profile for the first 24 hours and pharmacokinetic parameters of ivermectin are shown in figure 3 and Tables 1 and 2.

**Table 1).** Pharmacokinetic parameters of ivermectin alone after 1^st^ and 7^th^ dose described by non-compartmental analysis.

| PK Parameter | Units | Ivermectin 0.3 mg/kg |  |  |  | Ivermectin 1.2 mg/kg |  |  |  |
| --- | --- | --- | --- | --- | --- | --- | --- | --- | --- |
|  |  | 1<sup>st</sup> dose |  | 7<sup>th</sup> dose |  | 1<sup>st</sup> dose |  | 7<sup>th</sup> dose |  |
|  |  | Average | SD | Average | SD | Average | SD | Average | SD |
| AUC<sub>%Extrap</sub> | % | 29.2 | 4.1 | 17.6 | 11.5 | 37.4 | 17.8 | 19.1 | 19.1 |
| AUC<sub>24hr</sub> | hr*ng/ml | 2,152 | 460 | 6,481 | 853 | 10,188 | 2,781 | 28,495 | 7,190 |
| AUC<sub>INF</sub> | hr*ng/ml | n/a | n/a | 8,017 | 1,950 | n/a | n/a | 37,609 | 17,783 |
| Cl/F | L/hr/kg | 0.10 | 0.02 | 0.04 | 0.01 | 0.08 | 0.02 | 0.04 | 0.01 |
| Vz/F | L/kg | 1.90 | 0.41 | 1.03 | 0.25 | 1.73 | 0.61 | 1.15 | 0.16 |
| C<sub>max</sub> | ng/ml | 145.0 | 27.7 | 341.0 | 117.4 | 865.3 | 246.0 | 984.3 | 92.1 |
| C<sub>max</sub>/Dose | kg*ng/ml/mg | 483.3 | 92.4 | 1,136.7 | 391.3 | 721.1 | 205.0 | 820.3 | 76.8 |
| t<sub>1/2</sub> | hr | 13.1 | 1.9 | 19.2 | 6.9 | 16.5 | 6.0 | 24.1 | 8.3 |
| T<sub>max</sub> | hr | 3.3 | 1.2 | 6.7 | 4.6 | 2.7 | 1.2 | 5.3 | 2.3 |

**Table 2).** Pharmacokinetic parameters of ivermectin when co-administered with chloroquine after 1^st^ and 7^th^ dose described by non-compartmental analysis.

| PK Parameter | Units | Ivermectin 0.6 mg/kg + CQ 10 mg/kg |  |  |  | Ivermectin 1.2 mg/kg + CQ 10 mg/kg |  |  |  |
| --- | --- | --- | --- | --- | --- | --- | --- | --- | --- |
|  |  | First dose |  | Last dose |  | First dose |  | Last dose |  |
|  |  | Average | SD | Average | SD | Average | SD | Average | SD |
| AUC<sub>%Extrap</sub> | % | 24.8 | 6.4 | 1.3 | 1.1 | 30.4 | 13.5 | 4.5 | 5.7 |
| AUC<sub>24hr</sub> | hr*ng/ml | 6,742 | 415 | 18,333 | 8,989 | 10,406 | 2,793 | 38,079 | 41,331 |
| AUC<sub>INF</sub> | hr*ng/ml | n/a | n/a | 18,618 | 9,314 | n/a | n/a | 39,184 | 42,094 |
| Cl/F | L/hr/kg | 0.07 | 0.01 | 0.04 | 0.01 | 0.09 | 0.03 | 0.06 | 0.05 |
| Vz/F | L/kg | 1.10 | 0.19 | 1.20 | 0.29 | 1.55 | 0.47 | 1.60 | 0.75 |
| C<sub>max</sub> | ng/ml | 493.3 | 62.1 | 419.7 | 21.5 | 742.0 | 256.1 | 942.3 | 266.1 |
| C<sub>max</sub>/Dose | kg*ng/ml/mg | 822.2 | 103.5 | 699.4 | 35.8 | 618.3 | 213.4 | 785.3 | 221.7 |
| t<sub>1/2</sub> | hr | 11.5 | 2.5 | 25.3 | 12.1 | 13.3 | 4.5 | 28.3 | 23.6 |
| T<sub>max</sub> | hr | 4.0 | 3.5 | 4.0 | 0.0 | 4.0 | 3.5 | 4.0 | 0.0 |

**Figure 3).**
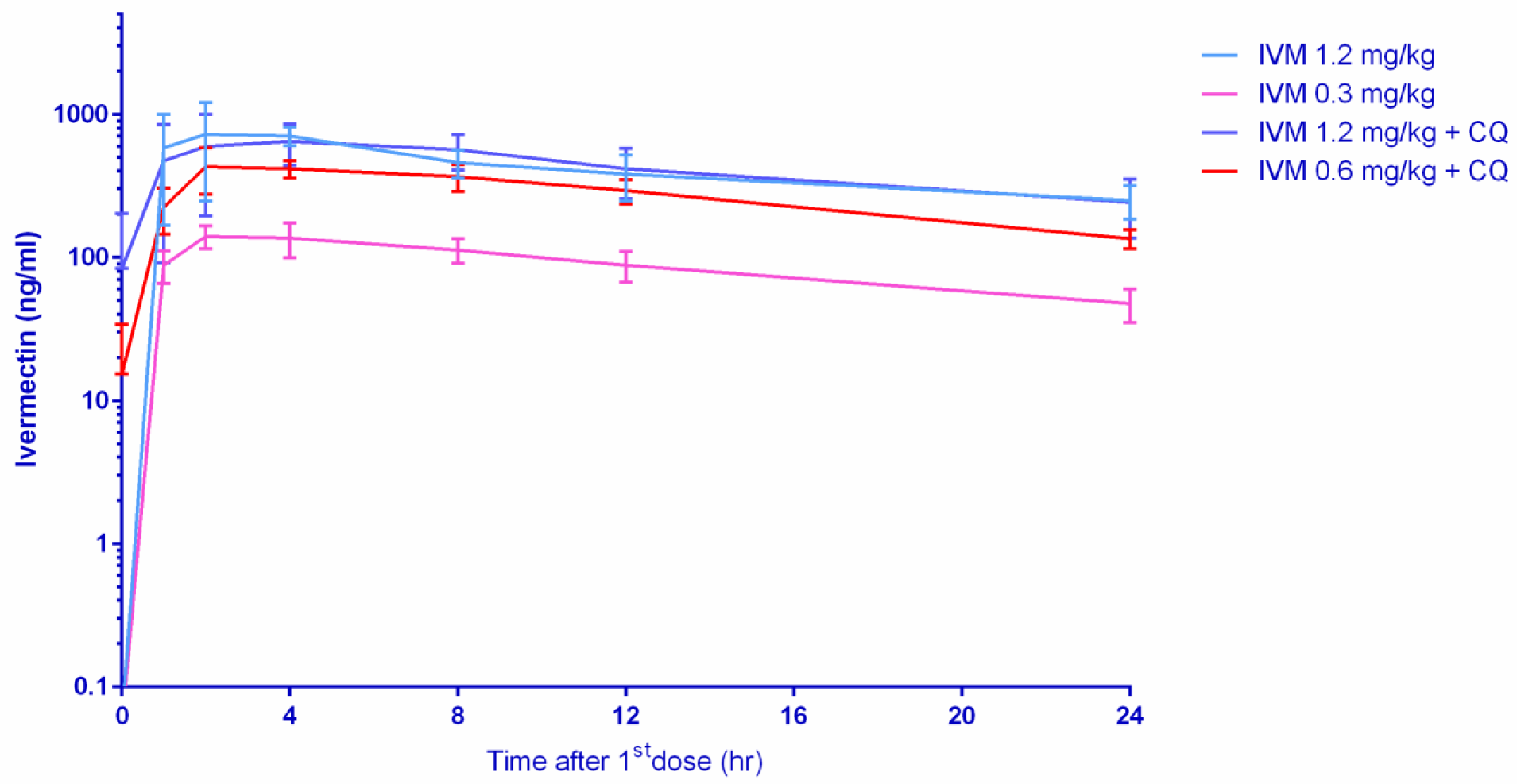
Ivermectin concentrations achieved in macaques 24 hours post first oral dose

Figure 3 represents the log concentration of ivermectin achieved in orally dosed macaques within 24 hours post the first dose. IVM = ivermectin, CQ = chloroquine (10 mg/kg).

Table 1 illustrates the pharmacokinetic parameters of ivermectin when administered alone after the first and seventh (last) doses. AUC_%Extrap_ is the percentage of area-under-the-curve infinity due to extrapolation from the last collection time point to infinity, AUC_24hr_ is the exposure through 24 hours, AUC_INF_ is the total exposure, Cl/F is the apparent clearance, Vz/F is the apparent volume of distribution, C_max_ is the maximum concentration, C_max_/Dose is the maximum concentration divided by the dose administered, t_1/2_ is the elimination half-life, and T_max_ is the time to reach the maximum concentration.

Table 2 illustrates the pharmacokinetic parameters of ivermectin when administered with chloroquine (10 mg/kg) after the first and seventh (last) doses. AUC_%Extrap_ is the percentage of area-under-the-curve infinity due to extrapolation from the last collection time point to infinity, AUC_24hr_ is the exposure through 24 hours, AUC_INF_ is the total exposure, Cl/F is the apparent clearance, Vz/F is the apparent volume of distribution, C_max_ is the maximum concentration, C_max_/Dose is the maximum concentration divided by the dose administered, t_1/2_ is the elimination half-life, and T_max_ is the time to reach the maximum concentration. CQ = chloroquine.

Figure 4 illustrates the linear pharmacokinetics of ivermectin as C_max_ and AUC increased in a dose dependent manner. Higher dose of ivermectin resulted in increased drug exposure with repeated dosing. Chloroquine did not interfere with ivermectin pharmacokinetics.

**Figure 4).**
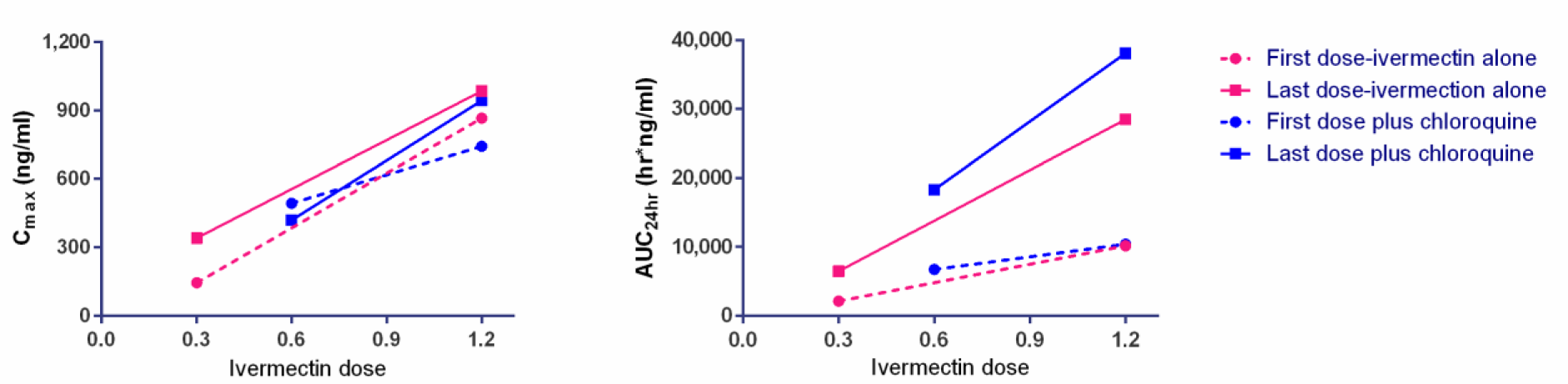
Relative ivermectin parameter values for C_max_ (left panel) and AUC_24 hr_ (right panel)

Figure 5 illustrates that ivermectin did not have any effect on chloroquine C_max_ or AUC_24hr_ (Paired Sample T-test P > 0.05). IVM = ivermectin, CQ = chloroquine, PQ = primaquine.

**Figure 5.**
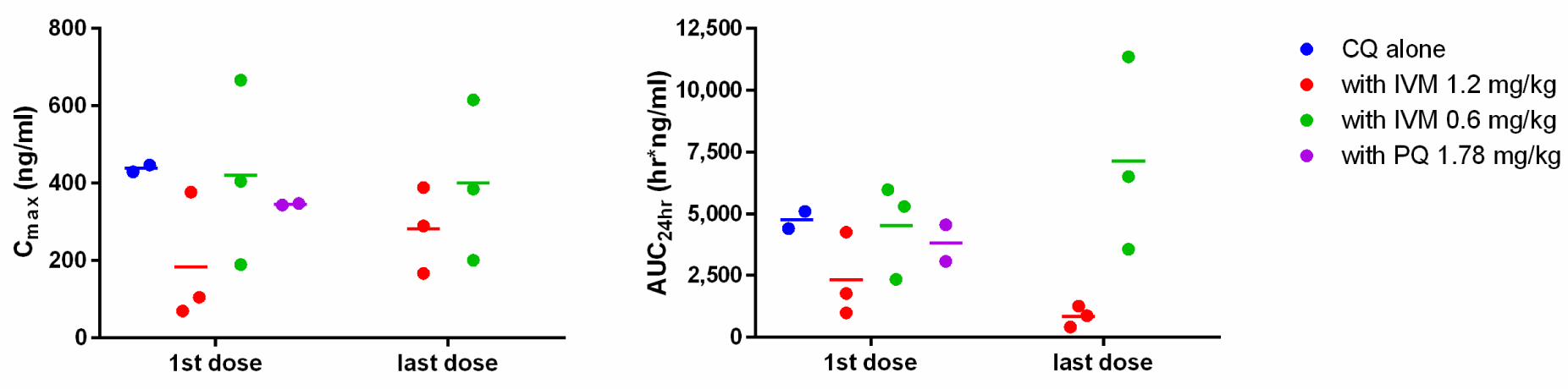
Relative chloroquine parameter values for C_max_ (left panel) and AUC (24 hours) (right panel)

Figure 6 illustrates the simulation of plasma ivermectin concentration-time profile. One-compartment analysis best described the observed data by using the estimates calculated by non-compartmental analysis following the first and seventh doses as initial estimates. In the simulation, C_max_ had mean estimates of 150, 300, and 600 ng/ml at approximately 4 hr post first dose and reached a steady state around the fifth dose with C_max_ at 243, 486, and 973 ng/ml at the ivermectin 0.3, 0.6, and 1.2 mg/kg dosing, respectively. IVM = ivermectin.

**Figure 6).**
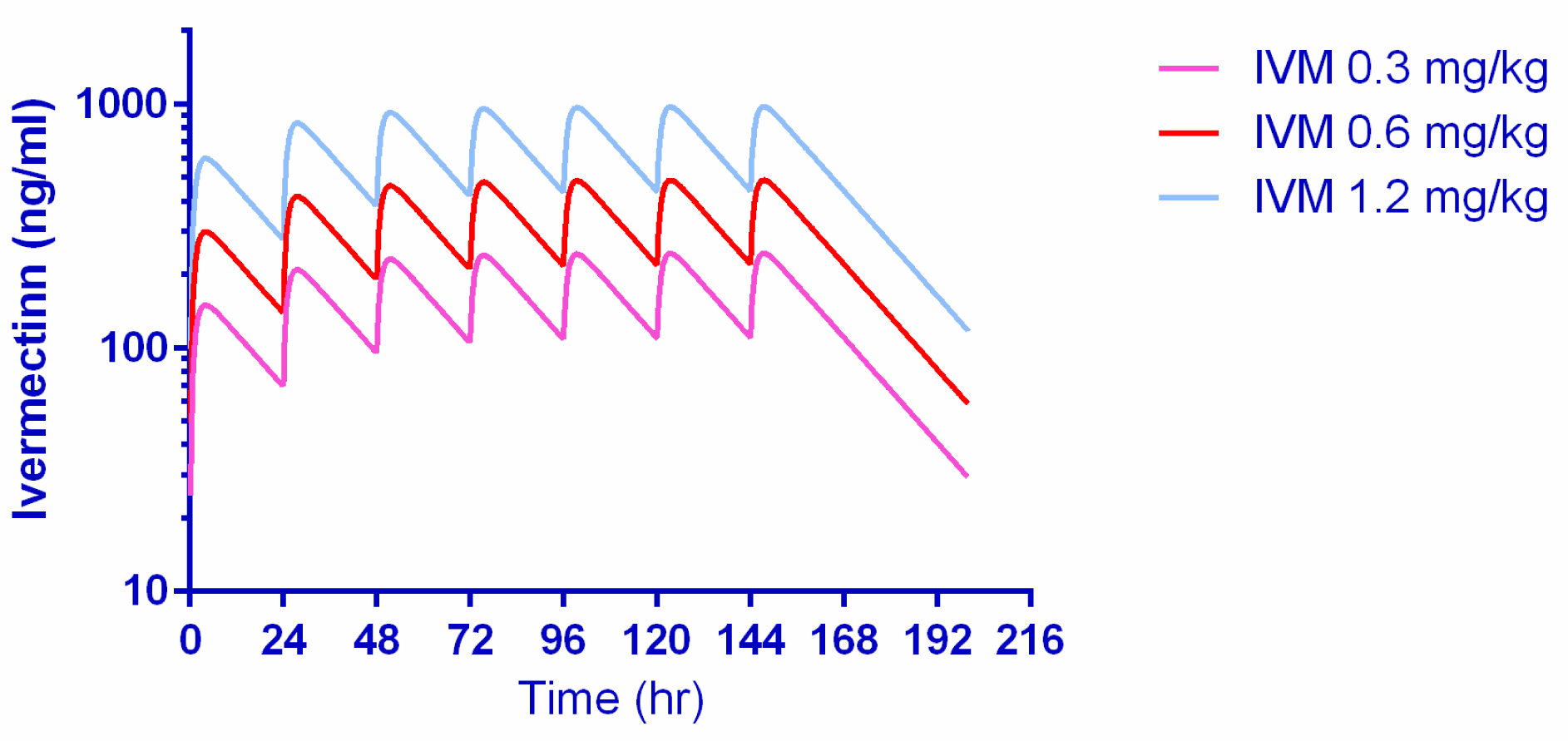
Pharmacokinetic simulation of ivermectin concentration-time profile when given as 0.3, 0.6 and 1.2 mg/kg for 7 days in Rhesus macaques

Figure 7 illustrates the mean ivermectin plasma concentration (ng/ml) by time (hr) profile 24 hours after the first and seventh dose with or without CQ (10 mg/kg). There was a slight reduction in peak concentrations achieved and delay in time to achieve peak concentrations when comparing the first and seventh doses. IVM = ivermectin, CQ = chloroquine.

**Figure 7).**
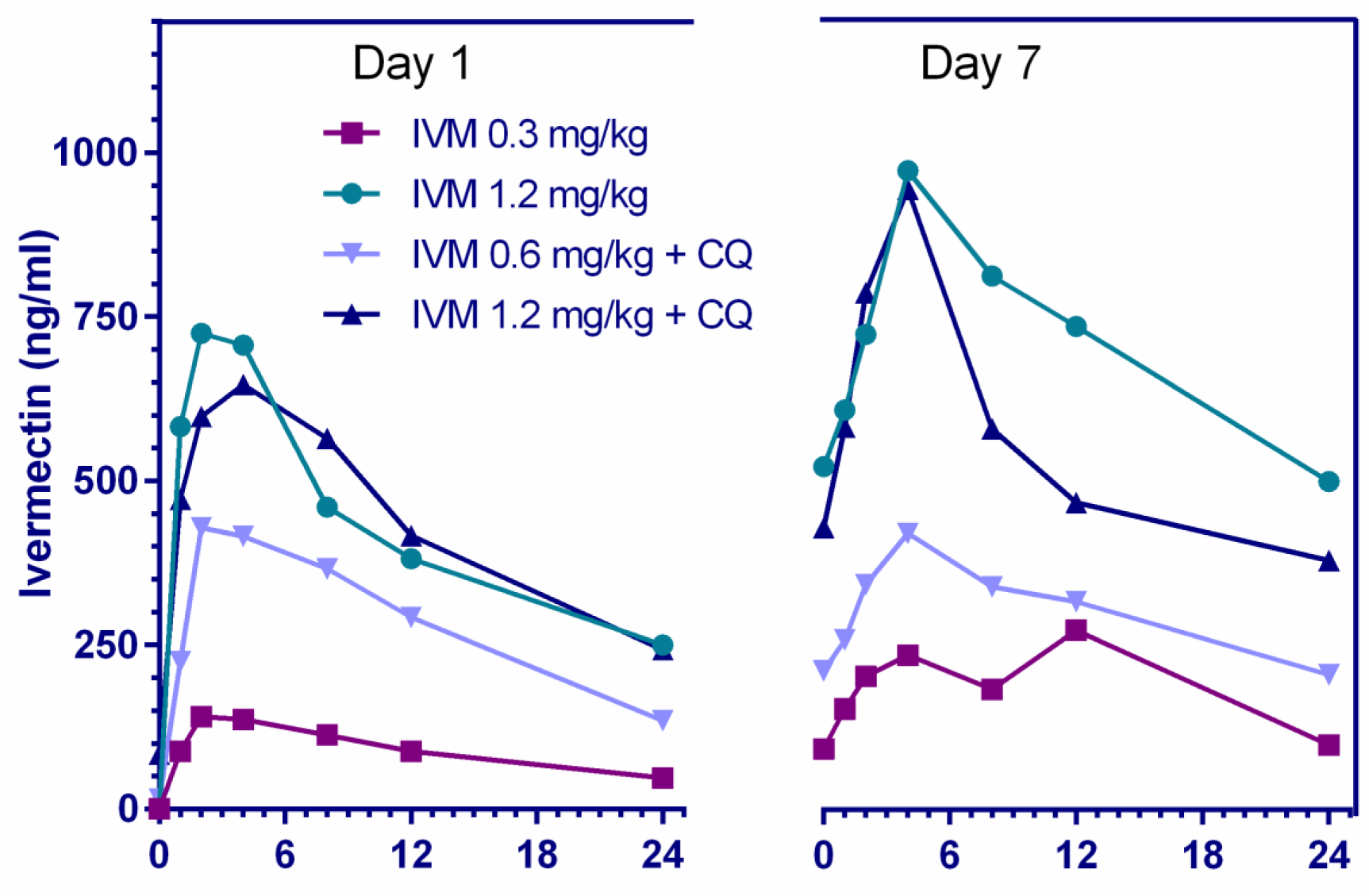
Mean plasma concentration-time profiles of ivermectin 24 hours after the first and seventh dose when administered ivermectin at 0.3, 0.6, and 1.2 mg/kg with and without chloroquine (10 mg/kg)

## Discussion

Ivermectin alone was safe and well-tolerated in macaques with repeated doses at 0.3 and 1.2 mg/kg for seven days, with no signs of neurological, gastroenterological, or hematological complications. One monkey vomited the first dose of ivermectin (1.2 mg/kg) when administered as monotherapy but had no emesis upon further dosing. Emesis was observed previously in ivermectin-treated macaques receiving 2 mg/kg single dose, and the occurrence of emesis increased with higher doses (4, 6, 8, 12, and 24 mg/kg) (17, 18). The combination of ivermectin (0.6 and 1.2 mg/kg) and chloroquine (10 mg/kg) for seven days was safe and well-tolerated in macaques.. This suggests that this combination could be used in humans during *P. vivax* MDAs in regions where chloroquine is still an effective *P. vivax* blood-stage therapeutic.

Prophylactic mode *in vitro* results with ivermectin parent compound indicate effect of ivermectin against *P. cynomolgi* liver schizonts and hypnozoites (Figure 1), but at higher concentrations than could be safely achieved in humans (10). However, there is a growing body of evidence that the activity of ivermectin is not restricted to the parent compound alone and that ivermectin metabolites may be active as well. Indeed, when comparing the effect of ivermectin metabolized by a human to that of parent compound mixed in human blood, the mosquito-lethal effect against *Anopheles dirus* and *Anopheles minimus* was 20- to 35-fold more potent (5) and the sporontocidal effect against *P. vivax* development in *An. aquasalis* was 5-fold lower (20). Even though *P. berghei in vitro* liver-stage IC_50_s were in the µg/ml range, liver schizont inhibition was achieved *in vivo* with ivermectin at doses plausible for use in humans (8). The points above warranted evaluation of ivermectin against *P. cynomolgi* in Rhesus macaques even though *in vitro* IC_50_s were in the µg/ml range and ivermectin only reaches ng/ml concentrations in orally-treated hosts.

There was no delay to patency of first blood-stage *P. cynomolgi* infection in either low- or high-dose ivermectin groups (Figure 2). Ivermectin displayed µM levels of liver schizont efficacy *in vitro*, however, a lack of delay to blood-stage patency suggests minimal impact of ivermectin on liver schizont development. Admittedly, the injection of one million *P. cynomolgi* sporozoites into the macaque sets a very high bar for any drug as it only requires one sporozoite to develop into a liver schizont to continue the blood-stage malaria infection. This is in contrast to a single mosquito that is predicted to deliver <100 sporozoites during blood feeding (21). The *in vitro* ivermectin experiments indicated prophylactic inhibition of *P. cynomolgi* hypnozoite development at µM concentrations, however, the macaque ivermectin challenge clearly demonstrated development of hypnozoites as indicated by the first and second blood-stage relapses occurring at approximately the same time as negative vehicle controls (Figure 2). Neither *in vitro* nor *in vivo P. cynomolgi* models indicate a radical cure efficacy potential for ivermectin. A recent human challenge trial (n = 8) with intravenous injection of cryopreserved *Plasmodium falciparum* sporozoites (n = 3,200) and a single oral dose ivermectin (400 µg/kg) failed to show liver-stage inhibition in terms of time to blood-stage patency (22).

To the best of our knowledge this is highest repeated dose ivermectin pharmacokinetic investigation in any mammal species. There were no significant changes in the Cl/F or T_1/2_. It should be noted that this study had a small sample size, only three macaques per ivermectin-treated group, and thus ivermectin autoinhibition warrants further evaluation in future trials. In humans, three repeated doses of ivermectin (30 or 60 mg) every third day did not inhibit C_max_ when comparing the first and third dose, suggesting a lack of autoinhibition (10). In FVB mice administered oral ivermectin (0.2 mg/kg) twice a week for five weeks there was a 1.7-fold reduction in 24 hour post-dose plasma ivermectin concentrations, while increasing the major metabolite concentration by 1.7-fold (23), suggesting induction of metabolism.

In macaques, co-administration of ivermectin (0.6 or 1.2 mg/kg) and chloroquine (10 mg/kg) for seven days was safe and well-tolerated. Co-administration of chloroquine and ivermectin did not have an effect on the C_max_ or AUC of ivermectin or chloroquine (Tables 1 and 2; Figure 5). The 1.2 and 0.6 mg/kg dose in macaques has an approximate HEDs of 0.55 mg/kg (total 3.85 mg/kg) and 0.27 mg/kg (total 1.89 mg/kg) respectively. This suggests that repeated daily dosing of ivermectin at 0.6 or 0.3 mg/kg could be used in combination with chloroquine in humans. While billions of ivermectin and chloroquine treatments have been administered to humans, there is very limited safety evidence for their co-administration. Only one study, on *Plasmodium vivax*, has co-administered ivermectin (0.2 mg/kg single-dose) and chloroquine (0.6 mg/kg first day, 0.45 mg/kg second and third day), in ten persons with no adverse events passively reported (20).

Ivermectin (24), chloroquine (25), and hydroxychloroquine (26, 27) have been shown *in vitro* to inhibit replication of the novel Severe Acute Respiratory Syndrome Coronavirus 2 (SARS-CoV-2). All three drugs distribute into lung tissues at higher concentrations than plasma for chloroquine and hydroxychloroquine in rats (28), for hydroxychloroquine in mice (29), and for ivermectin in goats (30) and cattle (31). Preliminary clinical evidence from an observational registry-based study found that COVID-19 patients that were ventilated and received single-dose ivermectin (150 µg/kg) had a significantly lower mortality rate, duration of hospitalization, and duration in the intensive care unit (32). The pre-clinical safety evidence in macaques presented here, *in vitro* efficacy, and preliminary clinical report on ivermectin efficacy in patients, warrants further investigation of ivermectin and chloroquine or hydroxychloroquine in SARS-CoV-2 infected persons.

This work verifies that the Rhesus macaque model provides a robust system for evaluating ivermectin pharmacokinetics. Newer formulations of ivermectin in development for humans, such as implants and expandable pill formulations (33, 34), could be evaluated in Rhesus macaques. Novel methods of *Plasmodium knowlesi* control, such as treatment of wild primates with ivermectin baits to target wild *Anopheles* populations could potentially be evaluated in this ivermectin macaque model system.

Although ivermectin was able to inhibit liver-stage development of *P. cynomolgi in vitro*, no demonstrable effect was observed with *in vivo* macaque challenge. Repeated doses of ivermectin (0.3, 0.6, 1.2 mg/kg) for seven days in macaques was safe with a corresponding rise in drug exposures (AUC), but no signs of autoinhibition. Co-administration of ivermectin (0.6 or 1.2 mg/kg) and chloroquine was safe and well-tolerated, with no drug-drug interactions altering ivermectin or chloroquine pharmacokinetics. Further ivermectin and chloroquine trials in humans are warranted for *P. vivax* control and SARS-CoV-2 chemoprophylaxis and treatment.

## Materials & Methods

### In vitro assay

The complete methodology is pending publication (A. Roth, personal communication). In brief, cryopreserved primary non-human primate hepatocytes (lot NGB) and hepatocyte culture medium (HCM) (InVitroGro^TM^ CP Medium) were obtained from BioIVT, Inc., (Baltimore, MD, USA) and thawed following manufacturer recommendations. The hepatocytes were plated into pre-collagen coated 384-well plates and used for experiments within 2 – 4 days after plating. Infectious sporozoites were obtained from *An. dirus* mosquitoes infected with *P. cynomolgi* B strain and used to infect the plated primary non-human primate hepatocytes. Ivermectin compound (Lot # MKBZ1802V, Sigma Aldrich, St. Louis, MO, USA) was dissolved in 100% DMSO and used at a final concentration of 100 µg/ml in an 8-point, 2-fold serial dilution. Ivermectin was administered in two treatment modes, prophylactic and radical cure. In prophylactic mode, drug was present for 4 days starting at point of sporozoite addition. Alternatively, in radical cure mode, drug was present for 4 days starting on day 4 post sporozoite inoculation.

Imaging and data analysis of the drug plates were completed using the Operetta CLS Imaging System and Harmony software 4.1 (Perkin Elmer, Waltham, MA, USA). Images were acquired using TRITC, DAPI, and bright field channels at 10x magnification. Parasites were counted with the TRITC channel and were identified by area, mean intensity, maximum intensity and cell roundness. Ivermectin IC_50_ curves and percent inhibition were generated using parasite population counts where controls were calculated as the average of replicates. The reported IC_50_s were obtained from two experimental replicates with three biological replicates for prophylactic mode and two biological replicates for radical cure mode, using an 8-point concentration format with 2-fold dilutions for final ivermectin concentrations of 0.781 to 100 µg/ml. The percent inhibition was performed using dose-response modeling in GraphPad Prism version 8.0 (GraphPad, La Jolla, CA, USA) where measured parasite quantity (hypnozoite or schizont parasites) were normalized to the negative control (infected wells) using the average of experimental and biological replicates.

### In vivo macaque trial

*Anopheles dirus* mosquitoes were used to produce *P. cynomolgi* (B strain) sporozoites, from a donor macaque infected with blood-stage *P. cynomolgi* parasites. For liver-stage challenge, each macaque was injected intravenously with 1 × 10^6^ *P. cynomolgi* sporozoites in a 1ml inoculum of PBS and 0.5% bovine serum albumin. USAMD-AFRIMS colony-born Rhesus macaques of Indian origin were used in this study. Ten healthy macaques, five male and five female, 3-5 years old and ranging in weight from 4.5-6.4 kg were selected for this study. All macaques were negative for simian retroviruses and simian herpes B virus. Two macaques served as negative controls and were treated initially with seven days of vehicle controls and treated with seven days of chloroquine (10 mg/kg) when parasitemia reached >5,000 parasites per µl at primary infection and first relapse, and with seven days chloroquine (10 mg/kg) plus primaquine (1.78 mg/kg) at second relapse. Two macaques served as positive causal prophylaxis controls and were treated initially with seven days of vehicle controls and treated with seven days of chloroquine (10 mg/kg) plus primaquine (1.78 mg/kg) at point of primary infection when parasites reached >5,000 parasites per µl. All study drugs were administered to restrained conscious macaques via nasogastric intubation at 1 ml/kg body weight.

Sparmectin-E (Sparhawk Laboratories, Inc., Lenexa, KS, USA) is a water-soluble formulation of ivermectin developed for oral use in horses. Ivermectin was diluted in sterile water and administered via nasogastric route. Six macaques received ivermectin; three low-dose (0.3 mg/kg) and three high-dose (1.2 mg/kg) for seven consecutive days starting one day before sporozoite challenge. If a primary blood-stage infection occurs, and blood-stage parasitemia reaches >5,000 parasites per µl, then the macaques receive seven days of chloroquine (10 mg/kg) plus ivermectin (1.2 mg/kg) for the high-dose group, and seven days of chloroquine (10 mg/kg) plus ivermectin (0.6 mg/kg) for the low-dose group. If a relapse occurs, and blood-stage parasitemia reaches >5,000 parasites per µl, then macaques received seven days of chloroquine (10mg/kg) plus ivermectin (1.2 mg/kg) for both the low- and high-dose groups. If a second relapse occurs, then the macaques were treated with seven days of chloroquine (10 mg/kg) and primaquine (1.78 mg/kg), terminating the experiment. Both the negative and positive control group macaques were treated with seven days of chloroquine (10 mg/kg) and primaquine (1.78 mg/kg) at the third relapse and first infection, respectively.

Macaques were observed several times in the first few hours post dosing, and at least three times a day for the remainder of the study for any clinical signs of neurological (*e.g.* ataxia, lethargy, imbalance) or gastroenterological (*e.g.* diarrhea, vomiting, weight loss) complications. Venous blood was collected at select time points and after macaques become blood smear positive for hematocrit, white and red blood cell count was determined.

### Parasitemia monitoring

#### Microscopy

Thick and thin blood smear samples were made and examined daily to quantify malaria parasitemia. Samples were fixed in methanol and stained with Giemsa stain. Blood smears were examined for the presence or absence of blood-stage parasites under oil-immersion objective. If no parasites were found in 50 microscopic oil-immersion thick fields or approximately 1,000 white blood cells (WBCs), the smear was considered negative. The parasitemia level was reported as number of parasites per 1µl or mm^3^ of whole blood. Parasites were counted per number of WBCs or red blood cells (RBCs) (i.e., per 1,000 WBCs or 1,000-10,000 RBCs). Parasitemia levels were calculated by the appropriate total blood cell count (white or red) per mm^3^.

#### Real Time PCR

Blood samples (0.2 ml) were collected on days 5, 6, and 7 after sporozoite injection. The same sampling schedule occurred in control macaques with the addition of sampling days 8, 9, and 10 (1ml) to obtain infected blood for controls used for method development. Blood was collected, stored in EDTA tubes, and kept frozen at −80°C. Parasite DNA was extracted from 200 ul from EDTA whole blood using EZ1 DNA blood kit with automated EZ1 Advanced XL purification system (Qiagen, Hilden, Germany). Real Time PCR for *P. cynomolgi* detection was performed by using Rotor Gene Q 5plex HRM platform (Qiagen, Hilden, Germany). Primer and probe were designed to target *P. cynomolgi* small subunit rRNA of blood-stage parasites (GenBank accession number L08242.1). Primer and probe sequences are as follows; *P. cynomolgi* Fwd: 5’-ATTGCGGTCGCAAATAATGAAG-3’, *P. cynomolgi* Rev: 5’-GGTATGATAAGCCAGGGA AGTG-3’, and probe: 5’ FAM-TACTCGCTCCTTCTGTTCCCTGGA-BHQ1-3’. Real Time PCR reaction was carried out in a total of 25 µl reaction using Rotor-Gene Multiplex PCR kit (Qiagen, Hilden, Germany) and a final concentration of primer and probe at 0.5 µm and 0.2 µm, respectively. PCR cycling condition consists of PCR initial activation step at 95°C for 5 mins followed by 45 cycles of denaturation at 95°C for 15 secs and annealing /extension at 60°C for 15 secs. The fluorescence data was acquired during annealing/extension step. Blood from a macaque (R915) previously infected with *P. cynomolgi* was used as a positive control and a cutoff at cycle 36 was used to define *P. cynomolgi* positive samples in this study.

### Pharmacokinetics

#### Sample collection and preparation

Blood sampling (1ml) for pharmacokinetic time points: just prior to first ivermectin dose, and after first dose 1, 2, 4, 8, 12 hours, then each consecutive day just before dosing, then after the 7^th^ dose at 1, 2, 4, 8, 12 hours, and days 1, 2, 5, 12, 19. If a primary infection occurred, then the same blood sampling schedule was repeated, but no blood for pharmacokinetics were collected at first or second relapses. Blood was collected in heparinized sodium Vacutainer tubes and centrifuged at 2,500 rpm for 20 min and then the supernatant (plasma) was transferred and kept at −80°C until analysis was performed. Plasma was separated into two tubes with 200-400 µl in each tube. Ivermectin was extracted using protein precipitation method by 2:1 of ACN (with IS):plasma volume, vortex mixed for 1 min and then centrifuged at 10,000 rpm for 10 min. 200 µl of supernatant fluid was filtered through a 0.22 µm PTFE membrane prior to inject to UPLC system.

#### Liquid chromatography-mass spectrometry analysis

The liquid chromatography-mass spectrometry (LC-MS) was performed on Waters Acquity UPLC^TM^ equipped with Waters Xevo® G2-XS QToF (Waters Corp., Milford, MA, USA). A Waters Acquity UPLC BEH C18 column (50×2.1 mm, 1.7 µm particle size) with precolumn of the same material was used to separate the compounds. The gradient mobile phase used for analysis of ivermectin was 5 mM ammonium formate and 0.1% formic acid in waters and methanol with the column temperature of 40°C, flow rate at 0.4 ml/min. The total run time was 7 min and the injection volume was 5 µl. For mass spectrometry was set in the positive electrospray ionization mode with multiple reaction monitoring. Instrument parameters included capillary voltage of 3.5 kv, source and desolvation temperature of 150 and 400°C, respectively. The nitrogen generator was set at 120 lb/in^2^ to generate cone and desolvation gas flow of 50 and 800 L/H, respectively. The mass transitions were observed at m/Z 892.77→569.50 and 894.79→571.52 for ivermectin and ivermectin-D2, respectively. Masslynx™ software (Waters Corp., Milford, MA, USA) was used for quantification.

#### Pharmacokinetic analysis

Noncompartmental analysis (NCA) was used to generate pharmacokinetic parameters using Phoenix WinNonlin 8.1 (Certara USA, Inc., NJ, USA). The PK parameters determined were the elimination half-life (T_1/2_), maximum concentration in plasma (C_max_), time to reach C_max_ after dosing (T_max_), area under the concentration-time curve in 24 hour (AUC_24hr_), area under the concentration-time curve after the last dose to infinity (AUC_INF_) and percentage of AUC_INF_ due to extrapolation from T_last_ (last collection time point) to infinity (AUC_%Extrap_) and since the fraction of dose absorbed cannot be estimated for extravascular models, apparent volume of distribution (Vz/F) and apparent clearance (CL/F) were substituted for V and CL. Data analysis and graphical representation were completed using GraphPad Prism version 8.0.

#### Pharmacokinetic modeling and simulation

Generated NCA pharmacokinetic parameters were used as parameters estimates for compartment modelling. Observed ivermectin concentration were best described by one compartment analysis with first order absorption and first order elimination.

### Ethical Statement

The USAMD-AFRIMS Institutional Animal Care and Use Committee and the Animal Use Review Division, U.S. Army Medical Research and Materiel Command, reviewed and approved this study (PN 16-03). Animals were maintained in accordance with established principles under the Guide for the Care and Use of Laboratory Animals eighth edition (35), the Animals for Scientific Purposes Act (36) and its subsequent regulations. The USAMD-AFRIMS animal care and use program has been accredited by the Association for Assessment and Accreditation for Laboratory Animal Care International (AAALACi).

## Acknowledgements

We thank the AFRIMS Department of Veterinary Medicine for conducting the macaque trial especially Laksanee Inamnuay, Kesara Chumpolkulwong, Natthasorn Komchareon, Chardchai Burom, Noppon Popruk, Sujitra Tayamun, Mana Saithasao, Alongkorn Hanrujirakomjorn, Nuttawat Wongpim, Khrongsak Saengpha, Phakorn Wilaisri, Chakkapat Detpattanan, Rachata Jecksaeng, Siwakorn Sirisrisopa, Yongyuth Kongkaew, Sakda Wosawanonkul, Sonchai Jansuwan, Amnart Andaeng, Chaisit Pornkhunviwat, Manas Kaewsurind, Wuthichai Puenchompu, Sawaeng Sripakdee, Dejmongkol Onchompoo, Thanaphon Rattanathan, Paitoon Hintong, Siwadol Samano and the Department of Entomology Malariology and Insectary Sections, especially Ratawan Ubalee and Siriporn Phasomkusolsil, for supporting the sporozoite production. Monoclonal antibody 7.2 (anti-GAPDH) was obtained from The European Malaria Reagent Repository (http://www.malariaresearch.eu). Funding sources include the Military Infectious Disease Research Program and the Defense Malaria Assistance Program. Funding support was provided by the University of South Florida College of Public Health Graduate Fellowship to A. Roth. The funders had no role in study design, data collection and interpretation, or the decision to submit the work for publication.

## Disclaimer

Material has been reviewed by the Walter Reed Army Institute of Research. There is no objection to its presentation and/or publication. The opinions or assertions contained herein are the private views of the author, and are not to be construed as official, or as reflecting true views of the Department of the Army or the Department of Defense. Research was conducted under an approved animal use protocol in an AAALACi accredited facility in compliance with the Animal Welfare Act and other federal statutes and regulations relating to animals and experiments involving animals and adheres to principles stated in the Guide for the Care and Use of Laboratory Animals, NRC Publication, 2011 edition.

